# Trehalose promotes wound healing *in vitro* by enhancing the migration of human keratinocytes via the VEGF/JNK/PI3K pathway

**DOI:** 10.1101/2025.08.25.672108

**Authors:** Keigo Taneda, Xiuju Dai, Kenji Watanabe, Teruko Tsuda, Hideki Mori, Ken Shiraishi, Yoichi Mizukami, Yasuhiro Fujisawa, Jun Muto

**Affiliations:** Department of Dermatology, Ehime University Graduate School of Medicine, Shitsukawa, Toon, Ehime, Japan; Institute of Gene Research, Yamaguchi University Science Research Center, Yamaguchi, Japan

**Keywords:** Keratinocytes, Trehalose, Vascular Endothelial Growth Factor A, Wound healing

## Abstract

**Background:** Trehalose is a naturally occurring disaccharide found in invertebrates but cannot be synthesized by vertebrates. We previously reported that high-concentration trehalose induces a transient senescent-like state in fibroblasts, leading to cell cycle arrest and growth factor secretion via CDKN1A/p21, and this process promoted keratinocyte proliferation, enhancing capillary formation and wound closure *in vivo*.

**Objective:** This study aimed to investigate the effect of trehalose on human keratinocytes.

**Methods:** Previously published RNA-seq data of cytokine-untreated samples from our group of trehalose-treated human keratinocytes were re-analyzed, and an *in vitro* scratch assay was performed using cells treated with mitomycin C.

**Results:** The trehalose-treated group exhibited increased wound closure. A significantly increased secretion of vascular endothelial growth factor (VEGF) was observed in keratinocytes treated with high-concentration trehalose, which is one of the most crucial molecules inducing angiogenesis in the skin. Significant upregulation of mRNA level and protein secretion of VEGF was confirmed using qPCR and ELISA, respectively. Furthermore, treatment with axitinib, a VEGF receptor inhibitor, significantly suppressed trehalose-induced activation of keratinocyte migration. Additionally, the increase in trehalose-induced migration activity was significantly inhibited by the Jun N-terminal kinase (JNK) inhibitor SP600125 and the PI3K inhibitor LY294002.

**Conclusion:** Trehalose promotes wound healing via VEGF secretion from keratinocytes and the PI3K and JNK pathways. The findings of this study may lead to the development of novel therapeutic agents that can alter the wound healing process.

**IMPORTANT:**

- Manuscripts submitted to Review Commons are peer reviewed in a journal-agnostic way.
- Upon transfer of the peer reviewed preprint to a journal, the referee reports will be available in full to the handling editor.
- The identity of the referees will NOT be communicated to the authors unless the reviewers choose to sign their report.
- The identity of the referee will be confidentially disclosed to any affiliate journals to which the manuscript is transferred.

**GUIDELINES:**

- For reviewers: https://www.reviewcommons.org/reviewers
- For authors: https://www.reviewcommons.org/authors

**CONTACT:** The Review Commons office can be contacted directly at

## 1. Introduction

Wound healing is a complex and dynamic process that requires the coordinated efforts of various cellular and molecular mechanisms to restore tissue integrity after injury. This important biological process usually involves four stages: hemostasis, inflammation, proliferation, and remodeling[1]. After injury, vascular permeability increases, and blood components are exuded, leading to platelets being concentrated at the wound site to seal the wound and repair the vascular damage[2]. The subsequent inflammatory phase triggers an immune response to protect against injury and infection. This phase is characterized by blood vessel expansion, increased blood flow, neutrophil and macrophage recruitment, and cytokine production. The next proliferative phase initiates tissue regeneration at the wound site. This phase is characterized by the progression of angiogenesis and the proliferation of fibroblasts around the wound, which produce collagen and extracellular matrix components for wound repair. At the same time, keratinocytes proliferate to form new skin that covers the wound surface [3]. The final remodeling phase involves remodeling and strengthening the tissue at the wound site. New blood vessels are no longer needed, and the blood supply to the wound area is reduced [1].

Successful wound healing requires a process called “re-epithelialization.” This process requires the directional migration of keratinocytes toward the wound center [4]. Chronic wounds or excessive scarring can occur when the wound healing process is impaired, posing significant clinical challenges. The c-Jun N-terminal kinase (JNK) pathway is a key signaling pathway involved in wound healing. JNK is a member of the mitogen-activated protein kinase (MAPK) family and plays a critical role in regulating cellular responses to stress, inflammation, and apoptosis [5]. The JNK pathway is involved in several key processes in wound healing, including regulating inflammatory responses, promoting keratinocyte migration, and re-epithelialization. JNK activation enhances keratinocyte migration, which is essential for covering the wound surface, and promotes fibroblast activity, which is crucial for extracellular matrix production and tissue remodeling [6].

Similarly, the phosphoinositide 3-kinase (PI3K) pathway plays a critical role in wound healing. PI3K is a family of lipid kinases that plays an important role in the regulation of various cellular functions, including cell growth, proliferation, survival, and migration. Upon PI3K activation by growth factors, cytokines, or other extracellular signals, it generates phosphatidylinositol (3,4,5)-trisphosphate, which serves as a docking site for proteins with pleckstrin homology domains, such as Akt (also known as protein kinase B) [7, 8]. Akt activation leads to the modulation of downstream targets that promote cell survival and growth. Therefore, the PI3K/Akt pathway plays a key role in tissue regeneration and repair. Recent studies have highlighted the importance of the PI3K pathway in wound healing and have shown that the downregulation of PTEN, a negative regulator of PI3K/Akt, promotes Akt activation and enhances wound healing [9]. The proper regulation of these pathways is essential for efficient wound healing and can be a potential therapeutic target, especially in treating chronic wounds.

Trehalose is a naturally occurring disaccharide consisting of two glucose molecules linked by an α,α-1,1-glycosidic bond. Trehalose is widely found in plants, fungi, bacteria, and some invertebrates and serves as a source of energy and a protective agent under stress conditions [10]. Trehalose is unique due to its remarkable stability and ability to protect cellular structures and proteins from damage caused by dehydration [11], freezing [12], and oxidative stress [13]. Furthermore, trehalose acts as a bioprotectant, which is one of its most remarkable properties [14]. During extreme environmental conditions such as drought and freezing, organisms that accumulate trehalose can stabilize their cellular membranes and proteins, allowing them to survive and recover once the conditions improve. This protective effect has led to the application of trehalose in various fields, including food preservation, pharmaceuticals, and cosmetics, where it enhances product stability and shelf life [15].

The beneficial effects of trehalose on skin health have been investigated. Trehalose is used in skincare products due to its moisturizing and antioxidant properties, which help protect and repair the skin barrier. Trehalose may have potential applications in wound healing and tissue regeneration due to its ability to stabilize proteins and cellular structures under stress conditions.

Generally, trehalose is a versatile molecule with wide-ranging applications from the food and cosmetic industries to potential therapeutic medical uses [16]. Given its unique properties, trehalose has become an important focus of ongoing research aimed at harnessing its full potential in various fields.

In our previous study, we reported the effects of high-concentration trehalose on dermal fibroblasts. Trehalose induced a transient senescent-like state in fibroblasts, leading to cell cycle arrest and growth factor secretion via CDKN1A/p21 [17]. This process promoted keratinocyte proliferation in living skin equivalent *in vitro*, enhancing capillary formation and wound closure *in vivo*. Therefore, this study aimed to investigate the effects of trehalose on human keratinocytes, specifically assessing changes in cell behaviors, such as cell proliferation and migration, after trehalose treatment. Additionally, this study aimed to investigate the effects of trehalose at the molecular level by examining growth factor secretion and key markers of the MAPK, JNK, and PI3K/Akt signaling pathways. The findings of this study may help develop new therapeutic agents that can alter the wound healing process.

## 2. Materials and methods

### 2.1. Keratinocyte culture and treatment

This study was approved in advance by the Ethics Committee of Ehime University School of Medicine (Ehime, Japan) and conducted in accordance with the principles of the Declaration of Helsinki. Written informed consent was obtained from all participants. Normal human skin biopsies were obtained from individuals undergoing plastic surgery. The epidermis was separated from the dermis, and normal human epidermal keratinocytes (NHKs) were isolated and cultured in a serum-free MCDB medium as previously described [18]. Cells were maintained in a humidified incubator at 37°C with 5% CO₂ and 95% air. Cells were preincubated in MCDB containing LY294002 (20 μM; Sigma-Aldrich, USA) or SP600125 (20 μM; Sigma-Aldrich) for 1 h before trehalose stimulation to inhibit the PI3K/AKT or JNK signaling pathways.

### 2.2. Preparation of RNA and real-time reverse transcription PCR

All probes specific for glyceraldehyde 3-phosphate dehydrogenase, vascular endothelial growth factor (VEGF), epiregulin (EREG), fibroblast growth factor (FGF2), and stem cell factor (SCF) were obtained from Thermo Fisher Scientific (Yokohama, Japan). Total RNA was isolated from NHKs and subjected to real-time reverse transcription PCR. Gene expression levels were analyzed as previously described [19, 20].

### 2.3. Scratch wound healing assay

After NHKs reached near confluence, the cell monolayers were scratched using a 200 μL micropipette tip, washed twice using phosphate-buffered saline, and incubated in an unsupplemented medium containing heparin-binding epidermal growth factor-like growth factor, with or without specific inhibitors. Phase-contrast images were captured at defined time points after scratching, and the percentage of the remaining wound area was calculated using ImageJ software (National Institutes of Health, Bethesda, MD, USA), relative to the initial wound area at 0 h (defined as 100%). Similar results were obtained in three independent experiments. In some experiments, cells were pretreated with LY294002 or SP600125 before scratching.

### 2.4. Growth factor quantification by LEGENDplex™ Multiplex Assay

The concentrations of multiple growth factors in the culture supernatants were measured using a bead-based multiplex immunoassay (LEGENDplex™ Mouse Growth Factor Panel, BioLegend, San Diego, CA, USA) according to the manufacturer’s instructions. Briefly, 25 μL of each sample or standard was mixed with 25 μL of premixed capture beads in a V-bottom 96-well plate and incubated for 2 h at room temperature with gentle shaking in the dark. After washing, 25 μL of detection antibodies were added and incubated for 1 h, followed by the addition of 25 μL of streptavidin-PE and an additional 30 min of incubation. The beads were then washed and resuspended in an assay buffer. Data were acquired using a BD FACSCanto™ II flow cytometer (BD Biosciences, San Jose, CA, USA). The results were analyzed using the LEGENDplex™ Data Analysis Software Suite (BioLegend).

### 2.5. Cell death assays

Cell viability was assessed using Cell Counting Kit-8 (Dojindo, Tokyo, Japan) according to the manufacturer’s instructions. The optical density was measured at 450 nm and was normalized to the corresponding stimulation control.

### 2.6. Whole-transcriptome analysis using RNA-seq

The RNA-seq data of cytokine-untreated samples from our previous study on trehalose-treated NHKs were re-analyzed [21]. Mapped read counts were normalized to transcripts per million, incremented by one across all values, and transformed into log2. Genes with a *P*-value <0.05 and a fold change >1.2 or <0.8 were selected and analyzed by Ingenuity Pathway Analysis (Qiagen). Heatmaps of upregulated and downregulated genes were generated based on transcripts per million values using Prism software (version 9.0; GraphPad Software).

### 2.7. Statistical analysis

At least three independent experiments were performed, all of which produced consistent results. Quantitative data were presented as dot plots using GraphPad Prism version 9.5.0 (GraphPad Software, San Diego, CA, USA). Each graph presents results from a single representative experiment, with 3–6 samples per condition. Individual dots represent the values of 3–6 replicates for each test point. The uncovered wound area was calculated using ImageJ version 1.53t (National Institutes of Health) and normalized to the wound area at 0 h, which was defined as 100%. Quantitative data were presented as mean ± standard deviation (SD), with n ≥ 3. Statistical comparisons were performed using Student’s *t*-test. Statistical significance was set at *P* < 0.05, *P* < 0.01, and *P* < 0.001.

## 3. Results

### 3.1. High-concentration trehalose promotes scratch wound closure in NHK layers through their ability to stimulate migration

A wound healing assay was performed under *in vitro* conditions to confirm the improvement of wound closure via enhanced migration by trehalose. An artificial wound was created on the NHK monolayer. Wound closure was observed 24 and 48 h after trehalose treatment with the addition of mitomycin C (Fig. 1A). Trehalose treatment increased wound closure by a +22% area ratio at 48 h compared with the untreated controls (Fig. 1B). High-concentration trehalose enhanced wound closure without promoting NHK proliferation, suggesting that its effect is due to increased cell migration (Supplementary Fig. 1). Trehalose up to 60 mg/mL did not affect cell viability, while concentrations above 100 mg/mL reduced it. Also, to assess whether the promotion of cell migration is specific to trehalose, a scratch assay with sucrose was conducted (Supplementary Fig. 2). Unlike trehalose, sucrose did not promote NHK migration at any concentration, indicating that the effect is specific to trehalose. These findings indicate that trehalose promotes NHK re-epithelialization by activating migration.

**Fig. 1.**
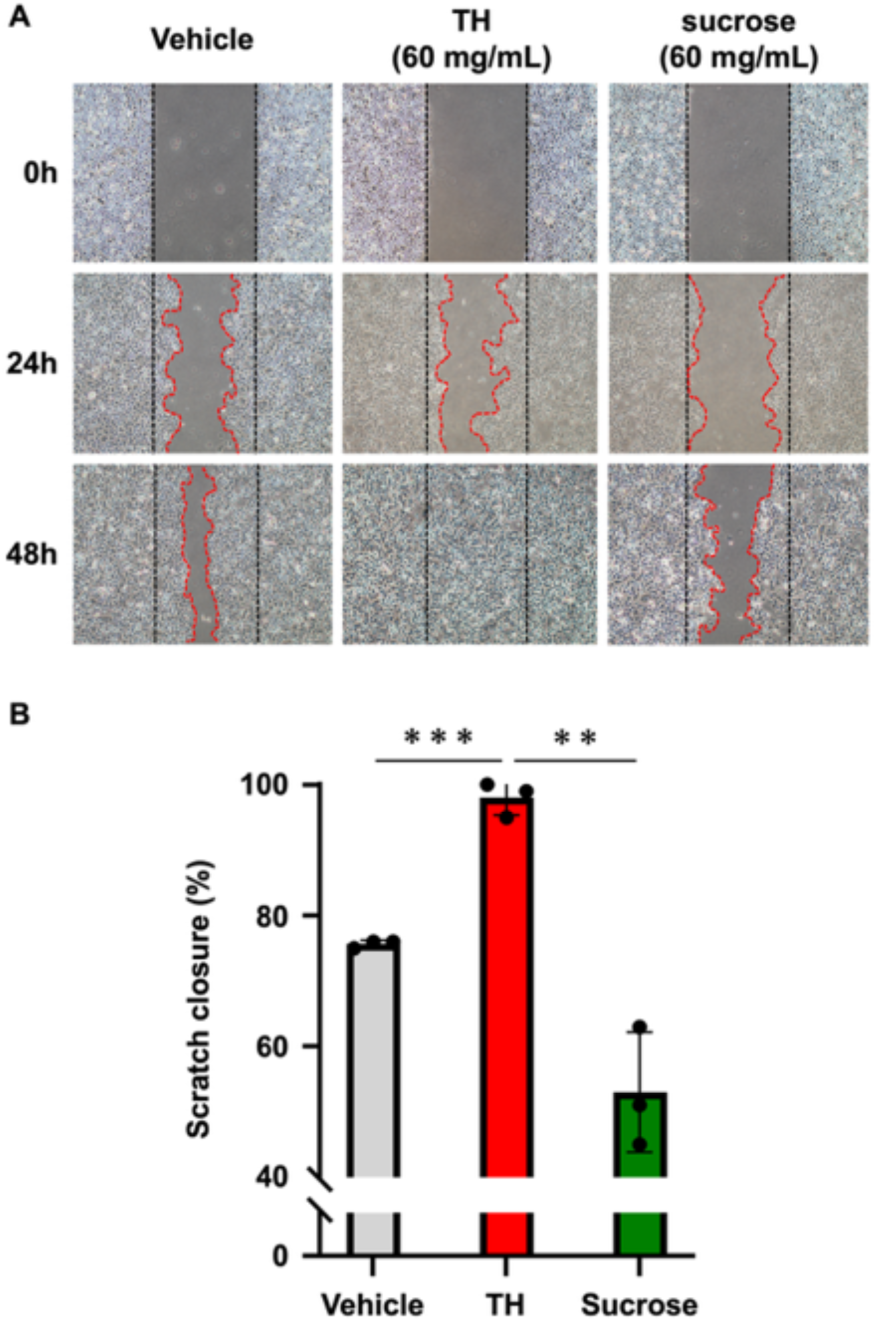
Scratch wound healing assay to determine whether trehalose promotes wound healing through cell proliferation or migration. (**A**) Micrographs from a phase-contrast microscope (10× magnification) show the results of the scratch wound healing assay with mitomycin C, capturing the healing rate of primary human keratinocyte monolayer at 0, 24, and 48 h after treatment with 60 mg/mL trehalose or sucrose or nontreatment. (**B**) Quantification of the area occupied by primary human keratinocytes after 48 h. Data are presented as mean ± standard deviation (SD) and are representative of three independent experiments. \**P* < 0.05, \*\**P* < 0.01, \*\*\**P* < 0.001, and \*\*\*\**P* < 0.0001 versus the vehicle-treated control group versus the sucrose-treated group by Student’s *t*-test.

### 3.2. Trehalose regulates several genes involved in cell migration

Previously published RNA-seq data from our group were re-analyzed to examine gene expression changes in trehalose-treated NHKs (60 mg/ml) to explore the mechanism that enhances NHK migration in the presence of trehalose. The dataset is available in GEO (accession number: GSE244738). Gene expression was significantly up- and downregulated by trehalose on a heatmap, indicating that trehalose affects various gene expressions related to skin formation (Fig. 2A). In the function analysis, the genes regulated by trehalose were associated with cell migration (Fig. 2B). Additionally, among the cell migration-related factors, VEGF was identified as a potential upstream factor (Fig. 2C).

**Fig. 2.**
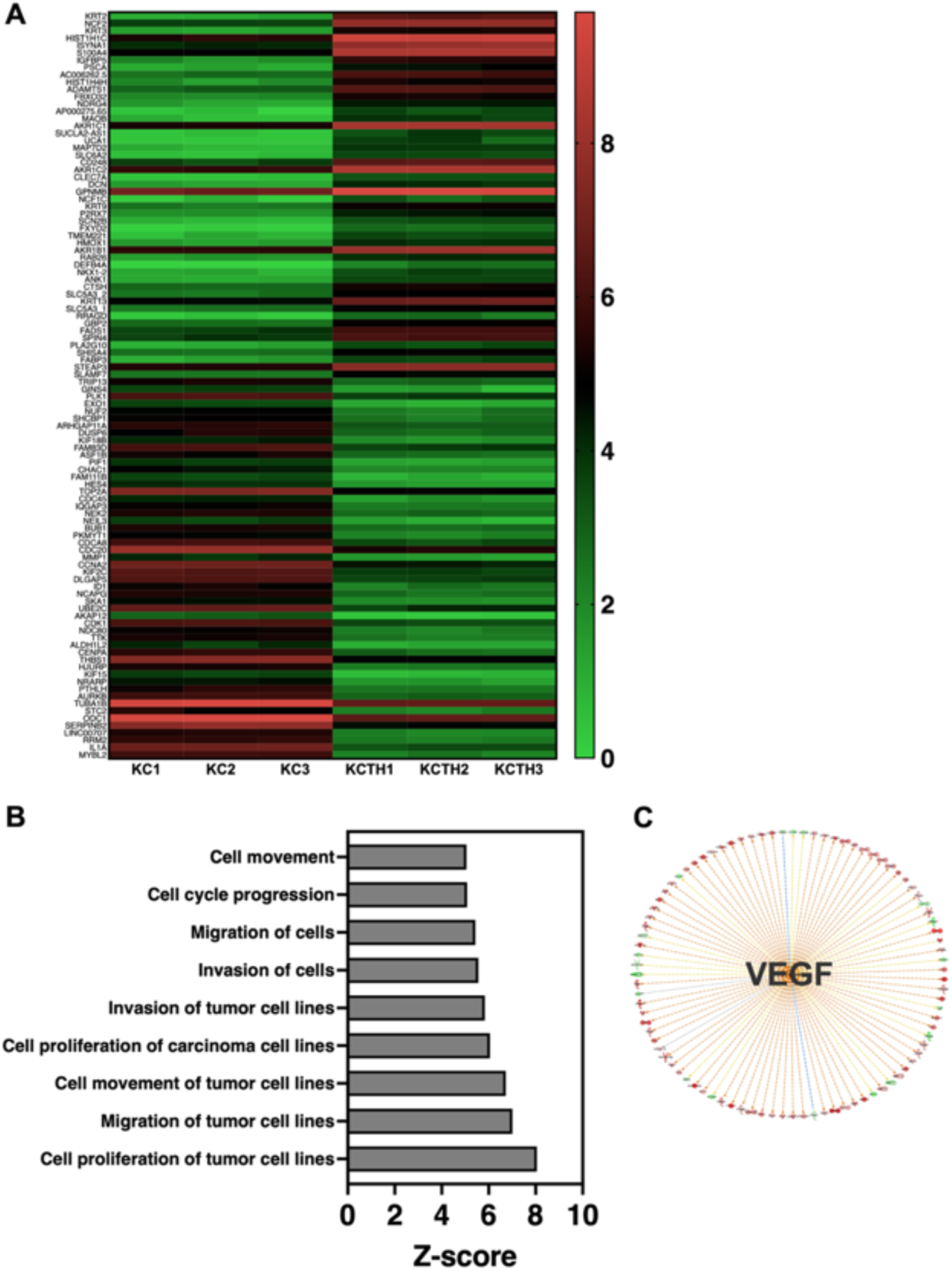
Trehalose-regulated gene expression in normal human epidermal keratinocytes (NHKs) analyzed by whole-transcriptome RNA-seq analysis. NHKs were treated with trehalose (60 mg/mL) for 18 h, and the total RNA was extracted and subjected to RNA-seq. (A) Heatmap representing the gene expressions significantly up- or downregulated by trehalose treatment based on transcripts per million. (B) Functional analysis revealed that trehalose-regulated genes are associated with cell migration. (C) Among the migration-related factors, VEGF was identified as a potential upstream regulator.

### 3.3. Trehalose induces an increase in the VEGF secretion from NHKs

Protein was quantified to elucidate the mechanism by which trehalose increases the migration activity. The culture supernatants of trehalose-treated NHKs were collected after 20 h. Protein concentrations in the supernatants were quantified in pg/mL using a bead-based multiplex LEGENDplex™ assay (LEGENDplex™ Custom Human Assay, Biolegend, San Diego, CA, USA) according to the manufacturer’s instructions. The results showed that trehalose had no effect on many proteins (Fig. 3A–C). Interestingly, NHK stimulation with trehalose significantly increased VEGF secretion (Fig. 3D). These findings indicate that VEGF plays a role in the enhanced migratory capacity induced by trehalose.

**Fig. 3.**
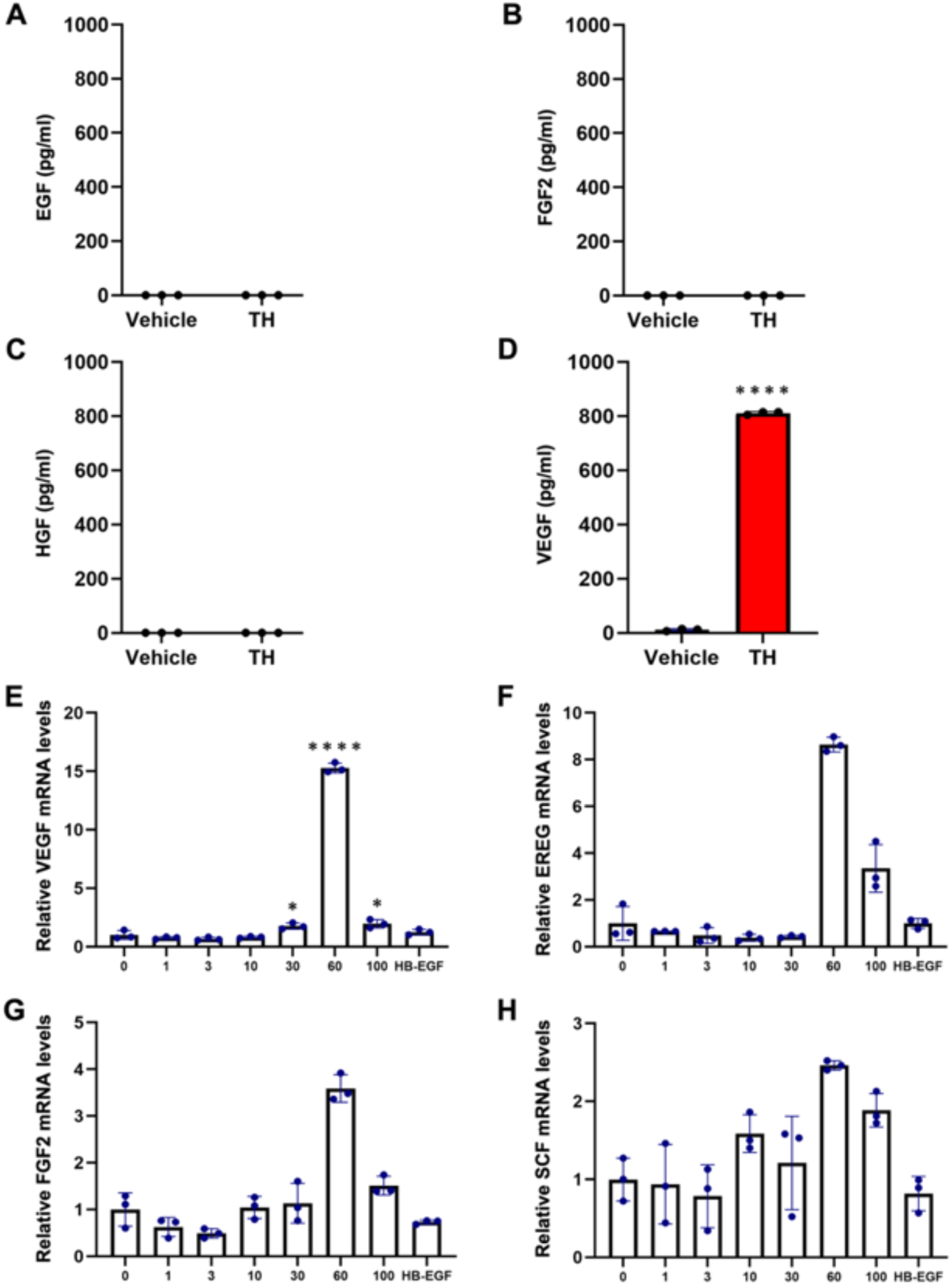
Trehalose modulates the expression of VEGF and growth factors of mRNA levels. Supernatants were collected and used for LEGENDplex™ assay to determine the levels of secreted: (**A**) EGF, (**B**) FGF2, (**C**) HGF, and (**D**) VEGF. (**E**) VEGF, (**F**) EREG, (**G**) FGF2, and (**H**) SCF mRNA expressions were assessed by qPCR. Data are presented as relative expression to control (vehicle-treated) primary human keratinocytes. Data are expressed as mean ± SD of triplicate wells and are representative of three independent experiments. \**P* < 0.05, \*\**P* < 0.01, \*\*\**P* < 0.001, and \*\*\*\**P* < 0.0001 versus the vehicle-treated control group versus the sucrose-treated group by Student’s *t*-test.

qPCR mRNA expression analysis of the wound healing-related genes was performed to confirm the results of the mRNA levels, which revealed that four genes (VEGF, EREG, FGF2, and SCF) significantly increased in NHKs treated with trehalose (60 mg/ml) for 24 h compared with vehicle control NHKs (Fig. 3E–H). FGF2 has been reported to promote NHK migration by activating Rac [22]. VEGF has also been reported to promote re-epidermalization and enhance wound closure [23]. In this way, the effect of trehalose on NHKs was confirmed even at the mRNA level.

### 3.4. VEGF promotes scratch wound closure in NHK layers through their ability to stimulate migration

A wound healing assay was performed under *in vitro* conditions to confirm that VEGF mediates improved wound closure via trehalose-promoted migration. NHKs were stimulated with trehalose and VEGF in the presence of mitomycin C for 24 h. After replacing the medium, an artificial wound was created on the NHK monolayer, and wound closure was observed at 48 h (Fig. 4A). VEGF treatment increased wound closure by a +22% area ratio at 48 h compared with the untreated controls (Fig. 4C). Furthermore, a comparison of VEGF and trehalose treatment showed no significant difference in wound closure, with only a −1% change after 48 h. These findings indicate that trehalose stimulates VEGF secretion, thereby activating migration and promoting NHK re-epithelialization.

**Fig. 4.**
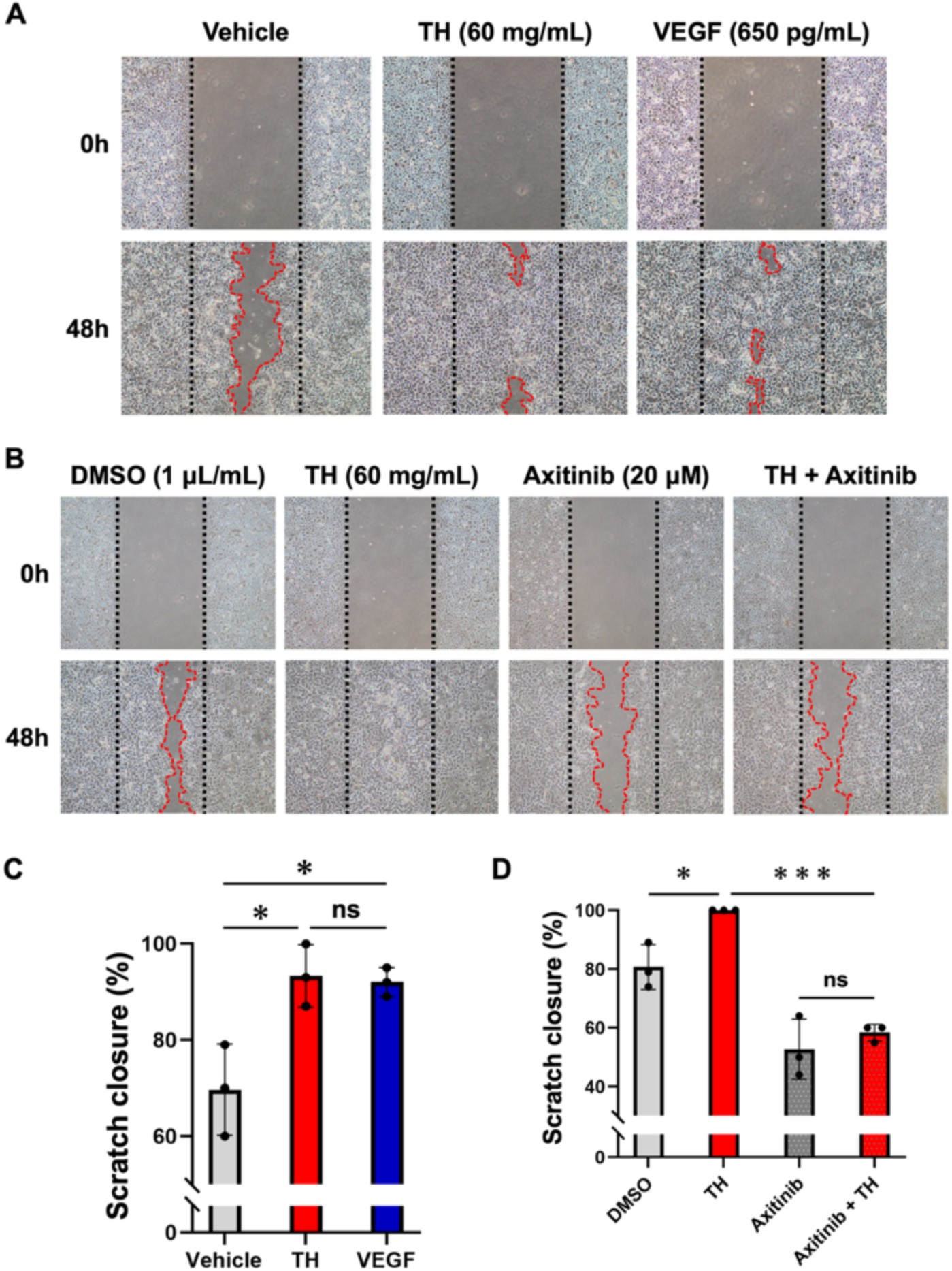
Scratch wound healing assay to determine whether VEGF promotes wound healing. (**A**) Micrographs from a phase-contrast microscope (10× magnification) show the results of the scratch wound healing assay with mitomycin C, capturing the healing rate of the NHK monolayer at 0 and 48 h after treatment with 60 mg/mL of trehalose, 650 pg/mL, or nontreatment. (**B**) Micrographs from a phase-contrast microscope (10 × magnification) of the scratch wound healing assay with axitinib and mitomycin C over the course of 48 h. (**C**) Quantification of the area occupied by NHKs after 48 h. Data are presented as mean ± SD and are representative of three independent experiments. ns: not significant. \**P* < 0.05 versus the vehicle-treated control group versus the trehalose-treated group versus the VEGF-treated group by Student’s *t*-test. (**D**) Quantitative analysis of percent closure of the scratch wounded areas of the NHK monolayer treated with trehalose with axitinib. Data are presented as mean ± SD and are representative of three independent experiments. \**P* < 0.05, \*\**P* < 0.01, \*\*\**P* < 0.001, and \*\*\*\**P* < 0.0001 versus the DMSO-treated control group versus the trehalose-treated group versus the axitinib-treated group versus the trehalose- and axitinib-treated group by Student’s *t*-test. DMSO was used as the negative control.

Furthermore, experiments were designed using a specific VEGFR inhibitor to determine whether trehalose promotes wound closure via VEGF production. After adding the VEGFR inhibitor axitinib (20 μM), scratch wound healing assays were performed on NHKs treated with trehalose for 24 h in the same manner as before (Fig. 4B). Similar to previous results, 60 mg/mL of trehalose dramatically promoted significant NHK migration after 48 h of incubation compared with the dimethyl sulfoxide (DMSO)-treated group (Fig. 4B and D). However, when combined with axitinib, the effect of trehalose in promoting wound closure was dramatically reduced (−42% area ratio). These findings indicate that VEGF plays a crucial role in the trehalose-induced enhancement of NHK migration.

### 3.5. Inhibition of growth and survival signaling pathways suppresses the migration-enhancing effects of trehalose

A scratch wound healing assay was performed using various inhibitors to elucidate how trehalose enhances the migration activity of NHKs. First, LY294002, a specific inhibitor of PI3K, was applied and experimented using the same protocol as before. Trehalose-induced NHK migration activity was suppressed in the presence of LY294002 (Fig. 5A and C).

**Fig. 5.**
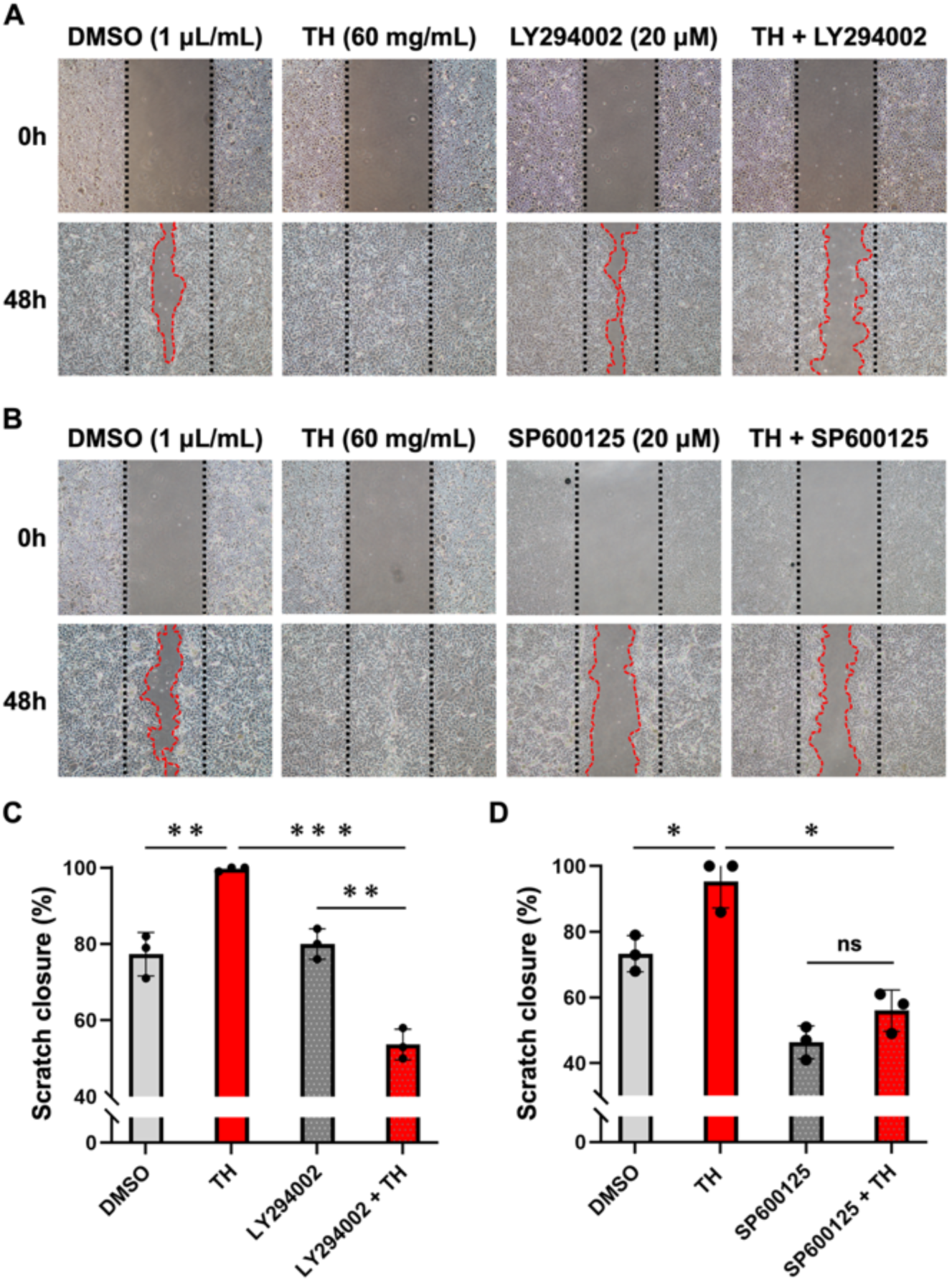
Scratch wound healing assay for determining the healing-promoting effects of trehalose in the presence of specific inhibitors of growth and survival signaling. (A, B) Micrographs from a phase-contrast microscope (10× magnification) of the scratch wound healing assay with LY294002 or SP600125 and mitomycin C over the course of 48 h. (C, D) Quantitative analysis of percent closure of the scratch wounded areas of the NHK monolayer treated with trehalose in the presence of LY294002 or SP600125. Data are presented as mean ± SD and are representative of three independent experiments. \**P* < 0.05, \*\**P* < 0.01, \*\*\**P* < 0.001, and \*\*\*\**P* < 0.0001 versus the DMSO-treated control group versus the trehalose-treated group versus the LY294002- or SP600125-treated group versus the trehalose- and LY294002- or SP600125-treated group by Student’s *t*-test. DMSO was used as the negative control.

Additionally, SP600125, a specific inhibitor of JNK, was applied, and experiments were performed using a similar protocol. SP600125 inhibited trehalose-induced NHK migration activity (Fig. 5B and D). These findings indicate that the effects of trehalose on enhancing cell migration activity are mediated through the PI3K and JNK pathways, which are known as growth and survival signaling pathways.

## 4. Discussion

Significant attention has been paid to trehalose due to its unique functions. Trehalose has been reported to have anti-inflammatory effects in femoral fractures [24], antiaging properties via anti-AGE activity [25], and the potential to improve diabetic symptoms through autophagy activation [26]. Furthermore, trehalose can promote significantly extensive spread of the epidermal layer of the living skin equiavalent [17] and enhance the barrier function of keratinocytes [21], with the former representing a groundbreaking discovery, demonstrating that high concentrations of trehalose can induce fibroblasts to enter a temporarily prohealing senescence-like state. A living skin equivalent exploiting this phenomenon may offer significant therapeutic potential for treating deep ulcers *in vivo*, which have historically been challenging to manage. The latter finding indicates that trehalose can restore the skin barrier by antagonizing IL-4/IL-13 signaling and suppressing STAT3/STAT6 activation *in vitro*. These findings indicate that topical application of trehalose is a promising therapeutic strategy for repairing skin barrier and preventing the onset of atopic dermatitis. Furthermore, this study showed that trehalose facilitates keratinocyte cell migration. This effect was mediated by the upregulation of VEGF, a key growth factor, which activates the JNK and PI3K signaling pathways. Notably, the inhibition of VEGF receptors and the JNK or PI3K pathways canceled trehalose-induced cell migration (Fig. 6). Interestingly, trehalose did not affect cell proliferation (Supplementary Fig. 1). Given the critical role of cell migration in wound healing [4], these findings indicate that trehalose positively contributes to skin wound repair. Furthermore, this effect was not observed when an equivalent concentration of sucrose (60 mg/mL) was administered (Fig. 1). These findings indicate that the observed enhancement of cell migration is not attributable to disaccharide-induced osmotic stress. Furthermore, no promotion of cell migration was observed at low or high concentrations of sucrose, indicating that this is an effect specific to trehalose (Supplementary Fig. 2).

**Fig. 6.**
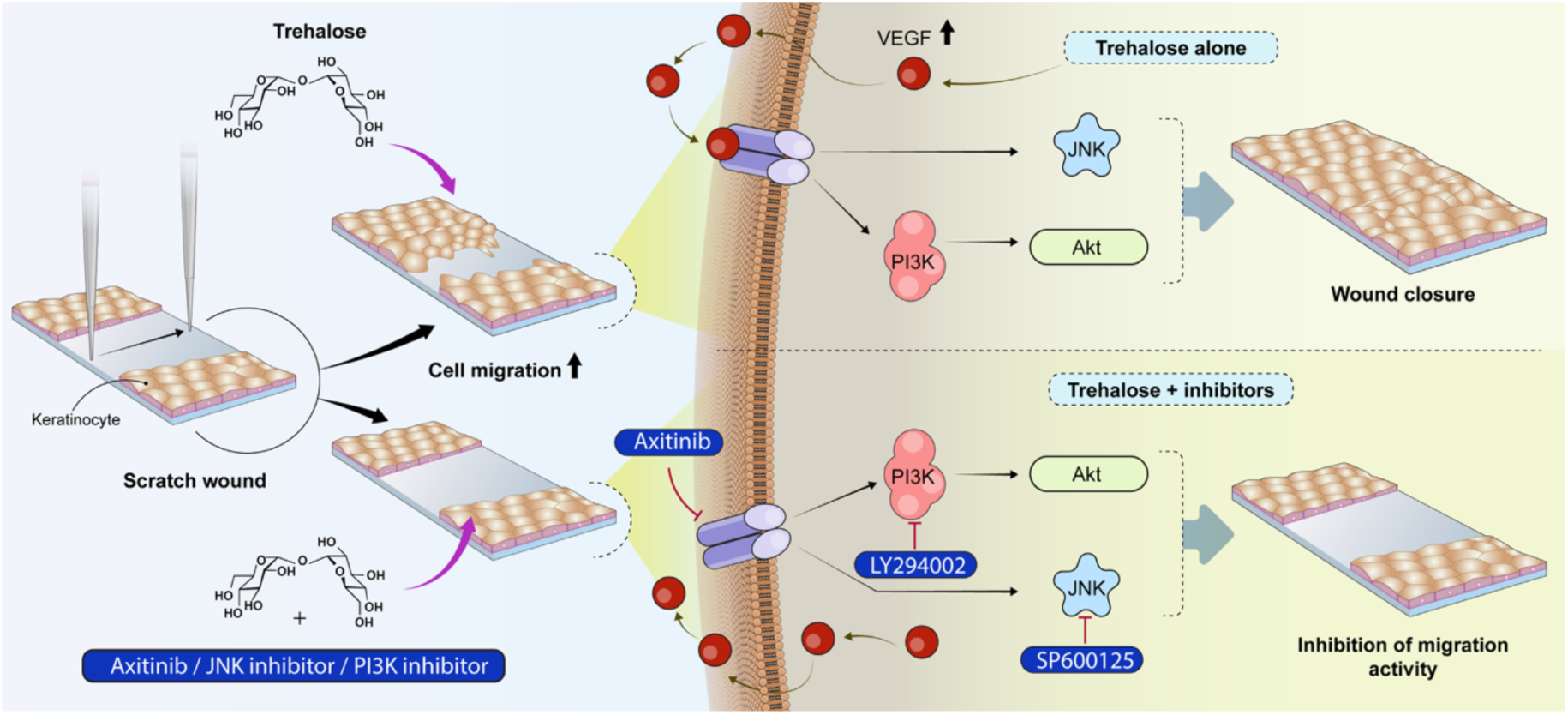
Putative signaling pathways involved in trehalose-induced wound repair in NHKs. Trehalose facilitates wound healing *in vitro* by the upregulation of VEGF expression and activation of the PI3K and JNK signaling pathways.

In this study, a series of experiments were conducted to elucidate the mechanism underlying the wound healing effects of trehalose. The findings indicate that trehalose enhances VEGF production. In mammals, the VEGF family comprises five members: VEGF-A, VEGF-B, VEGF-C, VEGF-D, and placental growth factor [27], with VEGF-A being the central and most widely studied member, commonly referred to as VEGF. Previous studies have shown that keratinocytes express all five VEGF receptors [28]. Therefore, VEGF secreted from keratinocytes upon trehalose treatment likely exerts its effects in a paracrine manner, thereby promoting cell migration. The expression of VEGF receptors in keratinocytes is crucial for maintaining skin homeostasis during wound healing. Several studies have shown that VEGF contributes to enhancing skin wound healing [27, 29]. These findings indicate that trehalose facilitates keratinocyte migration by promoting VEGF production and activating VEGF receptors.

Furthermore, various inhibitors were employed in this study to investigate the signaling pathways involved in trehalose-mediated wound healing. The JNK signaling pathway has been reported to be critical for wound healing by promoting keratinocyte migration [30]. Additionally, JNK has been reported to enhance the migration of keratinocytes by activating the PI3K/AKT and JNK pathways [31]. Therefore, we hypothesized that the JNK and PI3K/AKT pathways contribute to trehalose-induced wound healing. This study examined the effects of specific inhibitors to test this hypothesis. Trehalose-induced enhancement of keratinocyte migration was significantly inhibited in the presence of the PI3K inhibitor LY294002 and the JNK inhibitor SP600125. These findings indicate that VEGF production in keratinocytes activates the PI3K and JNK signaling pathways, thereby facilitating wound healing.

A key limitation of this study is the lack of in vivo evaluation; therefore, the wound-healing effects of trehalose under physiological conditions remain uncertain. Further studies using animal models, such as murine systems, are needed to validate our hypothesis and confirm the in vivo efficacy of trehalose.

In conclusion, this study showed that trehalose enhanced VEGF production and promoted wound healing *in vitro*. This effect was significantly suppressed by the VEGF receptor inhibitor axitinib and by inhibitors of the PI3K and JNK pathways. These findings indicate that trehalose facilitates wound healing by inducing VEGF release, thereby activating the downstream PI3K and JNK signaling pathways. Therefore, trehalose is a potential candidate compound for the development of novel therapeutic agents for skin wound healing.

## Declaration of interest

JM received research funding from ROHTO Pharmaceutical. The remaining authors state no conflict of interest.

## Funding

This work was supported by JSPS KAKENHI Grant Number JP24K11475 for Grant-in-Aid for Scientific Research (C).

## Declaration of generative AI and AI-assisted technologies in the writing process

During the preparation of this manuscript, the authors used the ChatGPT-4.0 model to assist in improving the clarity and accuracy of the language. All content was subsequently reviewed and edited by the authors, who take full responsibility for the final version. It is important to note that no part of the manuscript was generated directly by AI; the tool was used solely to refine the presentation of content originally written by the authors.

## Data availability statement

All data generated or analyzed during this study are included in this published article (and its supplementary information).

## Acknowledgements

We thank Eriko Tan for their technical assistance and thank Enago (www.enago.jp) for the manuscript review and editing support.

## CRediT authorship contribution statement

Conceptualization: JM;

Data Curation: KT, XD, YM, KW, JM;

Formal Analysis: KT, XD, YM, KW, JM;

Funding Acquisition: JM;

Investigation: KT, XD, YM, KW, TT, JM;

Methodology: KT, XD, YM, KW, JM;

Project Administration: KT, XD, JM;

Resources: JM;

Visualization: KT, XD, YM, KW, JM;

Writing e Original Draft Preparation: KT, XD, YM, JM;

Writing e Review and Editing: KT, XD, YM, KW, KS, HM, YF, JM

## Supplementary information

### Effects of trehalose on NHK cell viability

Cell proliferation rates were measured using Cell Counting Kit-8 (Dojindo, Tokyo, Japan) to determine whether the acceleration of wound closure by high-concentration trehalose was due to increased cell proliferation or migration. The results showed that 60 mg/mL (and below) trehalose did not significantly affect the cell viability of NHKs cultured in a serum-free medium compared with the untreated group. However, trehalose concentrations above 100 mg/mL significantly reduced NHK cell viability (Supplementary Fig. 1). Additionally, no significant difference in cell viability was observed in cells treated with sucrose, a disaccharide similar to trehalose. These findings indicate that trehalose can enhance wound closure through its ability to stimulate NHK migration.

### Effects of sucrose on NHK cell migration

To confirm whether the promotion of cell migration by disaccharides is specific to trehalose, a scratch assay using sucrose was performed. An artificial wound was created on the NHK monolayer. Wound closure was observed 24 and 48 h after scurose treatment with the addition of mitomycin C (Supplementary Fig. 2). The addition of sucrose, regardless of concentration, did not enhance cell migration.

**Supplementary Fig. 1.**
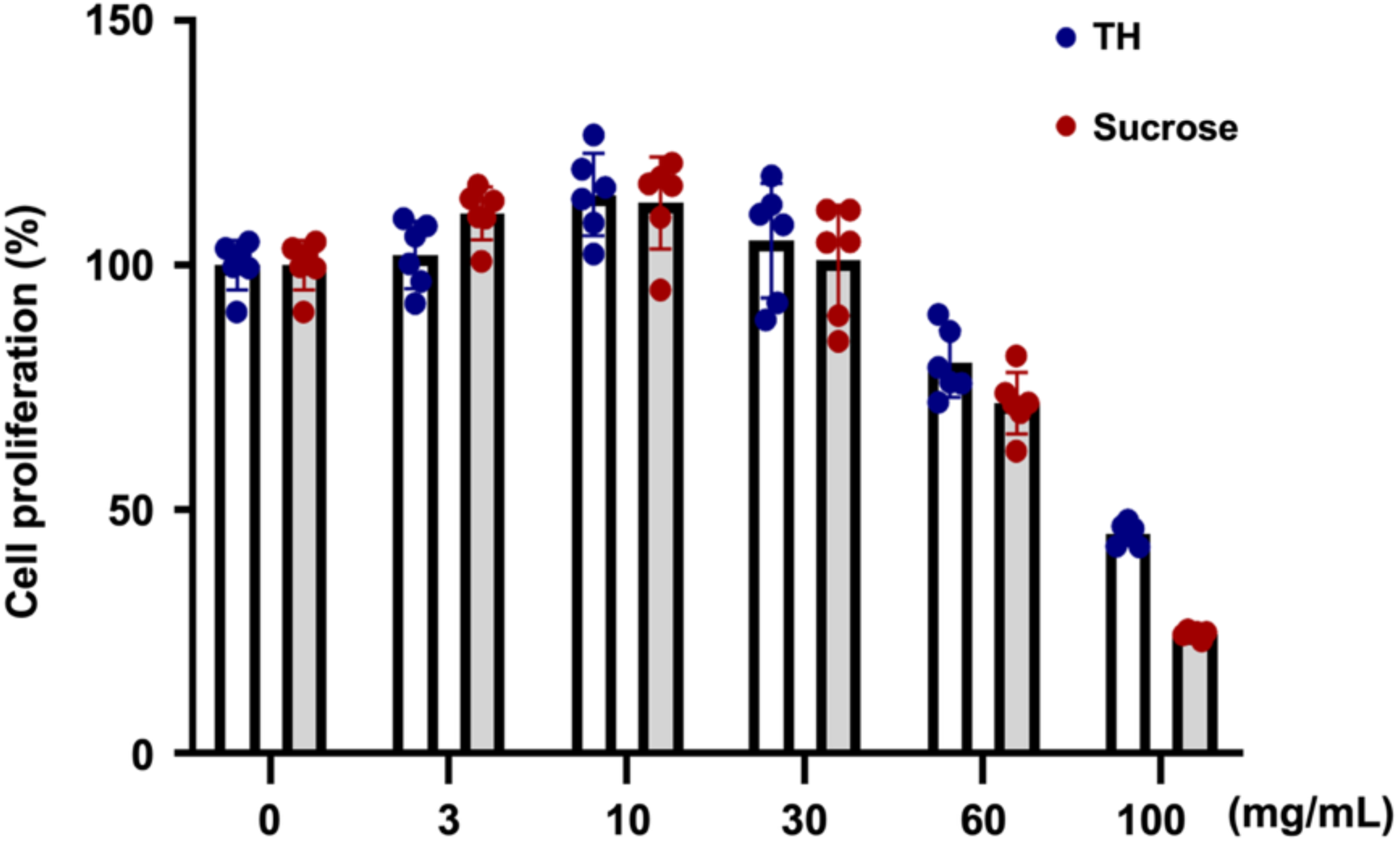
Cell viability test in NHKs with various concentrations of trehalose for 24h. Cell viability was determined and expressed as a percentage of the control (without trehalose treatment). Data are presented as mean ± SD (n = 6).

**Supplementary Fig. 2.**
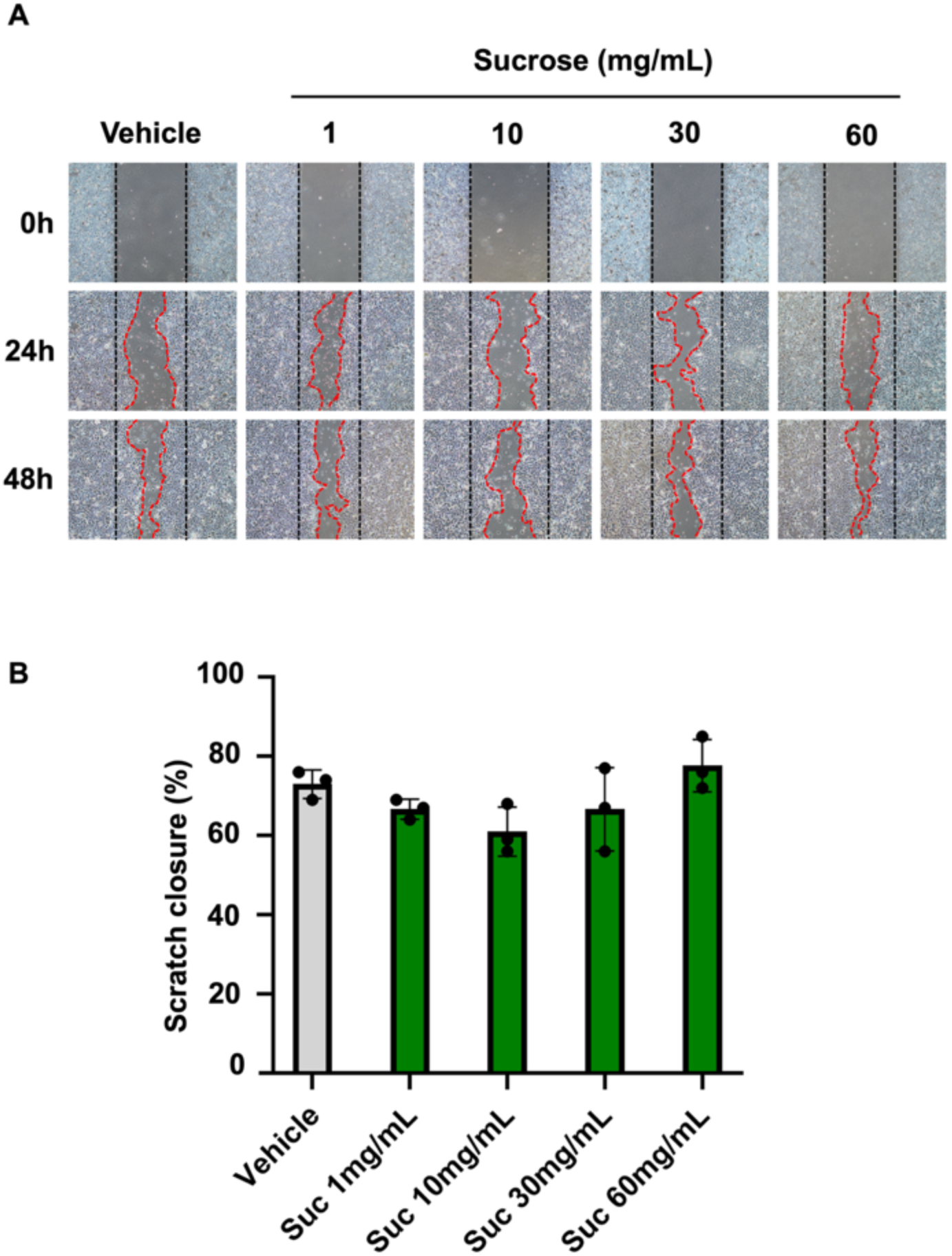
Effect of sucrose on scratch wound closure in NHKs monolayers. (A) Micrographs from a phase contrast microscope (10× magnification, scale bar =100 μm) capturing cell monolayer healing rate of NHKs at 0, 24 and 48 h after treatment with 1, 3, 10, 30 and 60 mg/mL of sucrose or untreated. (B) Quantification of the area occupied by NHKs after 48 h. Data were expressed as means ± SD and are representative of three independent.

